# What is the storage effect, why should it occur in cancers, and how can it inform cancer therapy?

**DOI:** 10.1101/2020.03.28.013557

**Authors:** Anna K. Miller, Joel S. Brown, David Basanta, Nancy Huntly

## Abstract

Intratumor heterogeneity is a feature of cancer that is associated with progression, treatment resistance, and recurrence. However, the mechanisms that allow diverse cancer cell lineages to coexist remain poorly understood. The storage effect is a coexistence mechanism that has been proposed to explain the diversity of a variety of ecological communities, including coral reef fish, plankton and desert annual plants. Three ingredients are required for there to be a storage effect: 1) temporal variability in the environment, 2) buffered population growth, and 3) species-specific environmental responses. Here, we argue that these conditions are observed in cancers and that it is likely that the storage effect contributes to intratumor diversity. Data that show the temporal variation within the tumor microenvironment is needed to quantify how cancer cells respond to fluctuations in the tumor microenvironment and what impact this has on interactions among cancer cell types. The presence of a storage effect within a patient’s tumors could have substantial impact on how we understand and treat cancer.

## Introduction

Ecosystems in nature include coexisting species that compete for space, resources, and safety from predators^1–4^. Similarly, tumors exhibit micro-environmental heterogeneity that contains coexisting cancer cell types that experience variations in nutrients, toxic metabolites, and diverse types of normal cells^5^. In nature, mechanisms of coexistence can explain the diversity of species within a community. Typical mechanisms include food-safety tradeoffs (one species is the better competitor while the other is better at avoiding predation), diet separation (each species has subset of resources on which it is the more successful consumer), habitat selection (each species has a habitat within which it is more successful), and competition-colonization trade-offs (one species slowly outcompetes the other at a given spot while the other is more successful at dispersing to unoccupied spots). Similar mechanisms likely explain some of the diversity of cancer cell types within and among a patient’s tumors^6^.

Long underappreciated in ecology was a mechanism of coexistence now known as the *storage effect*^*7–11*^. Three ingredients can promote the coexistence of species by the storage effect: 1) temporal variability in environmental conditions that include periods favorable and unfavorable for survival and reproduction (we shall refer to these as good and bad), 2) the presence of two life history states within each species (one conducive to proliferation during good periods, the other conducive to survival during bad periods), and 3) differences in how species perceive and invest effort to grow, or do not, in good and bad periods, reflecting some underlying trade-offs and adaptation to somewhat different environmental conditions.

Here, we posit that the storage effect may promote the coexistence of at least some cancer cell types within the tumors of some or many different cancers. To demonstrate the plausibility of the storage effect we will discuss how, as in ecosystems in nature, conditions (1) and (2) are universal properties of most tumor microenvironments. Next, we discuss why there may be tradeoffs in the way cancer cells experience good and bad periods. We then discuss how the storage effect manifests in nature, followed by a simple mathematical model for cancer tumor cells, illustrating how the storage effect works. We conclude with discussion of how knowledge of the presence of a storage effect in a patient’s disease might inform therapies.

## Temporal Variation in Nature and in the Tumor Microenvironment

Very few, if any, ecological communities experience temporally constant environments. Migratory birds escape the harsh winter conditions of higher latitudes by migrating to destinations closer to the equator. Deciduous trees lose their leaves during dry and/or cold seasons. Year to year variation in temperature and precipitation may portend droughts or floods and hot spells and cold snaps. Fire, disease, or pestilence occur episodically within ecosystems. A beaver damming a stream can raise water tables, drown surrounding terrestrial vegetation and create a wetland. This wetland can revert when the beaver dam breaks or decays. Virtually all living organisms experience predictable and unpredictable variability in environmental conditions that influence growth, reproduction, and survivorship.

Like natural ecosystems, cancer cells inhabiting a tumor microenvironment experience both regular and irregular temporal fluctuations. Blood flow changes due to unstable vasculature on timescales of minutes to hours^12^. This instability arises as cancer cells adapt to local hypoxic conditions by recruiting new blood vessels, which enable the delivery of nutrients and the removal of toxic metabolites so that cancer cells can survive when nutrients are low. However, as cancer cells induce the growth of new blood vessels, the vascular network becomes more irregular and disorganized leading to unpredictable spatial and temporal variations in blood flow^13^. Consequently, transient changes in perfusion lead to cycling hypoxia. Local fluctuations between hypoxia and reoxygenation affect cells adjacent to the poorly perfused blood vessels. This is in contrast to chronic hypoxia, which affects cells far away from vessels due to limited diffusion. Cycling hypoxia is associated with an increase in cell migration, metastatic potential, and resistance to treatments compared to chronic hypoxia^14^.

While changing patterns of blood flow likely create the greatest source of temporal variability in oxygen, pH, immune infiltration, nutrients and toxins, the architecture of cells within and around a tumor microenvironment also may change temporally. The rate of diffusion of nutrients (such as glucose and glutamine) towards a neighborhood of cancer cells, and the diffusion of metabolites away (such as lactate, free-oxygen radicals and pyruvate) may vary temporally as other neighborhoods of cancer cells block or unblock intracellular channels, as immune cells move in or out of an area, and as fibroblasts change inter-cellular matrices and/or the secretion of growth factors. Regardless of its source or regularity, coexisting cancer cells experience considerable temporal variability in opportunities and hazards that likely create good and bad periods in terms of their proliferation and survival rates, setting the potential for the storage effect to promote diversity of cancer cells within tumors.

## Proliferative and Non-proliferative Life History States in Nature and in Cancer

In response to fluctuations that generate good and bad periods, many organisms have evolved different life history states; one that is best at exploiting the good times and the other best at surviving the bad times. Many single-celled protists have a proliferative state that allows the cell to feed, move and proliferate^15^. This state however may not be able to survive bad periods when their pool of water dries up or the environment offers only toxins and no nutrients. Such protists also have an encysted state. This state, while unable to proliferate, is highly resistant to desiccation, toxins and nutrient deprivation. This state may be the only way the protist survives through bad periods. Having a fraction of the protist population in an encysted state means that some opportunities are lost during good periods, but survival is insured during the bad. Baker’s yeast is a classic example. The dry packet of encysted yeast can survive for years without opportunity for growth and reproduction. Upon activation and favorable conditions these yeast enter a feeding and proliferative state. Among species in nature, “quiescence” or “dormancy” refers to a range of cellular or organismal states characterized by slowed metabolism and relatively high resistance to hardships from their surrounding environment^16–21^.

In cancer, cells that are reversibly arrested in the G0 phase of the cell cycle are referred to as quiescent, whereas “cell dormancy” typically refers to a long-term quiescence^22^. Many cancer cells respond to hypoxia by becoming quiescent and upregulating autophagy to survive lower levels of nutrients and oxygen^13^. Cancer cells may survive harsh environments (e.g., hypoxia, low pH, toxins) by forming poly-aneuploid cancer cells (PACCs) (also referred to in the literature as polyploid giant cancer cells (PGCCs)), a population of reversibly quiescent cells that form rapidly (within 72 hours) in response to environmental stress and later divide asymmetrically to repopulate a tumor with cells of normal ploidy that have increased resistance to chemotherapy^23,24^. In their non-proliferative state, dormant or quiescent tumor cells are able to evade treatments that preferentially target highly proliferative cells. After treatment, these cells can resume proliferation which can result in tumor growth and relapse. Quiescent states are ubiquitous in cancer and can be associated with metastasis and relapse of cancerous growth^25^. As such, dormancy and quiescence challenge our ability to treat, control, or moderate cancer.

In nature, quiescent and resistant states can have a powerful effect. They can allow populations to survive despite exposure to conditions that limit or preclude population growth. They can buffer populations against variation in favorability for growth on many time scales, including seasonal, annual, and multiannual; they can contribute to more diverse communities than would exist without them^9,10,26–28^. Given how common dormancy/quiescence is in cancer, we expect to find similar ecological effects of these life-history stages for the cancer ecosystem. Quiescent or dormant states could maintain diverse cancer phenotypes that may proliferate, coexist, or displace each other over the course of time within the patient. For instance, disseminated cancer cells that stay dormant, sometimes for many years, and then reemerge as metabolically active cells that proliferate, often leading to death of patients. These emergent metastases appear to result from enhanced cell lineage persistence through dormancy^23–25,29,30^. The movement of cells into and out of dormancy, as the environment around them changes, could affect what clones co-occur in a tumor or in a patient by changing the ways in which cell-cell interactions such as competition play out at the level of populations of different cell types. The presence of quiescent and dormant, resistant states of cancer cells sets the potential for the storage effect to promote diversity of cancer cells within tumors. These possibilities have clear relevance for cancer control.

## Evidence for Population-Specific Behaviors and Trade-offs in the Way Cancer Cells Experience Good and Bad Times

As in nature, where a year with plentiful but not excessive water would be expected to be generally a good year for vegetation, cancer cells also would share many environmental needs and so experience good and bad times in part together. However, as with plants that differ in tolerance to low water availability or high temperatures, cancer cells differ in their sensitivities to hypoxia, low pH, immune infiltration, absence of growth factors, and the stromal (created by normal cells) architecture of their microenvironment, generating potential for trade-offs that make some periods better for some cell clones and others better for other clones. The heterogeneity of cancer cells in many features, including their proliferative potential, hormone receptor expression, immunogenicity, sensitivity to drugs, motility, and angiogenic potential^31^ enables cancer cells to have *population-specific environmental responses*, an essential component of the storage effect. For example, in stage 2 invasive breast cancers, cells on the tumor edge tend to have an acid-producing invasive, proliferative phenotype whereas those in the hypoxic tumor core have a less proliferative phenotype^32^. Tumors also have metabolic heterogeneity in the form of acid-resistant or glycolytic phenotypes^33^. Mathematical models of cancer evolution suggest that high nutrient variability (cycling hypoxia) gives a competitive advantage to cancer cells that have a higher rate of phenotypic transition, which promotes high levels of phenotypic heterogeneity^34^.

Cancer cells have a wide range of behaviors associated with entering and exiting from non-proliferative states, which provide the *buffering life history stage* that enables a storage effect. Cancer cells may remain in these states for times ranging from the very short (one to several days) to the very long (decades, as observed for emergence of metastatic cancer growth after years of healthy remission)^25,35^. In fibroblasts, cells move deeper into quiescence following longer durations of serum starvation or contact inhibition and require a stronger growth stimulation to exit quiescence compared to cells in shallow quiescence^36^. Time of exit from quiescence exit can vary within a clonal population, even when cultured in the same growth conditions^37^. Dormant cells awaken due to environmental changes. Bone is highly dynamic, undergoing constant remodeling, which can reawaken dormant cancer cells ^38^. The growth of new blood vessels provokes sufficient change to the microenvironment to reactivate breast cancer cells^39,40^. Age-related inflammation promotes the outgrowth of dormant cells in pancreatic ductal adenocarcinoma liver metastases^41^. PACCS emerge from quiescence after removal of chemotherapy and they do so after a range of quiescence durations (Sarah Amend, PhD, email communication, February 2020). Drug resistance is associated with a heterogeneous population of non-proliferative cells, including drug-tolerant persisters, therapy-induced senescent cells, hypoxic drug resistant, and disseminated tumor cells. These populations differ in their propensities to remain in or exit the growth arrested state^22^.

## A Simple Model of the Storage Effect in Cancer

Mathematical models have been useful to understand cancer biology and to help test what treatment strategies might improve patient outcomes over standard of care. Many model types have been used, including game theory models^42–49^, agent based models (ABMs)^33,50–55^, and Lotka-Volterra type dynamical models that consider cancer clonal cells as interacting populations^56,57^. So far, very few of these models consider variation in the environment of cancer cells in time. It is unsurprising that temporal variation in the tumor environment has had little attention in dynamical models. This was long the case in models of natural ecosystems, and the data available for cancer tend to be static snapshots of a tumor. Radiographic and histologic images show cancer at a point in time. Although they reveal, and provide data for, modeling spatial heterogeneity in cancer, they do not capture or inform the potential variation in time that small-scale physiological studies indicate is there as well ^12,14,58^.

Stochastic dynamic models of interacting populations of cancer cells are a straight-forward way to incorporate temporal environmental variation and are the class of models that have primarily been used to develop understanding of the storage effect. The simple Lotka-Volterra type population models so far used to study cancer are not sufficient to capture coexistence mechanisms such as the storage effect because they do not incorporate a non-proliferative life history state nor do they incorporate species-specific demographic responses to a varying environment^3,59,60^. Variation in the environment can be proxied as stochastic variation in demographic parameters, such as entry to and exit from quiescence and rates of growth, reproduction, or survival if cancer cells.

To show how coexistence can occur via the storage effect we construct a simple discrete-time stochastic model that includes two species, each with two life history stages (proliferative and quiescent), and stochastic variation in good and bad periods.

### Illustrating the Storage Effect

The plausibility of the storage effect promoting the coexistence of different cell types within a tumor microenvironment emerges from 1) temporal variability in blood flow, immune infiltration, stromal architecture, and physical conditions, and 2) the widespread use of cell arrest, quiescence, and/or dormancy that allows cancer cells to exist in two states – proliferative and non-proliferative. Coexistence can occur if there is a tradeoff in how two cell types experience good and bad periods.

To illustrate how storage effects can work we propose a simple model that aims to keep assumptions to a minimum while containing the essential elements. We imagine that each cancer cell type has two stages: an arrested state, and a proliferative state. A fraction of cells, θ, leave the arrested state during each period. Those that enter the proliferative state suffer a mortality rate that depends on whether it is during a “good” or “bad” period. These periods could represent a microenvironment of a tumor where fluctuations in blood flow, hypoxia and/or growth factors contribute to times of plenty and times of famine. Upon emerging from the arrested state, cells successfully survive to proliferate at fraction s. Let *s*_*g*_ >> *s*_*b*_ be these survival rates during good and bad periods, respectively.

The biology here is that if a cell attempts to be proliferative it may lack key resources even as the cell commits to cell division. The consequence is cell mortality that would not have happened had the cell remained in an arrested state. For example, providing estrogen to breast cancer cells that otherwise have no other nutrients can cause mortality, as cells are tricked into trying to proliferate (Robert Gatenby, MD, email communication, February 2020). Those cells that survive produce a maximum number of daughter cells, *f*, that declines with competition from other cells in a proliferative state. We use a Ricker model to describe the competition between proliferative cells for resources. In the absence of competitors, a proliferative cell produces its maximum number of daughter cells; with competitors, the production of daughter cells declines exponentially with the density of other proliferative cells. We imagine that *f* can take on a positive value that represents the expected number of daughter cells that might accrue during a period sufficiently long to permit one or more rounds of cell division. At the end of the period, daughter cells enter the pool of cells in an arrested state.

Now let there be two cell types that are equal in all ways except that they do not always experience good and bad periods in the same way. While unlikely that two cell types would be identical, this assumption allows us to illustrate coexistence that only manifests because of the storage effect.

For cancer cell types, it is likely that what is good for one is, in part, good for the other type. However, differences in nutrient metabolism, sensitivity to growth factors, resistance to hypoxia, ability for immune evasion, and/or physical conditions like pH means that at times one cell type may experience a bad period while the other a good period. Let *q* be the probability of a good period, and let 0 ≤ *r* ≤ 1 represent the degree to which the two cell types experience good and bad periods similarly. The term (1−*r*) then represents the probability that the two cell types are experiencing the type of period independently of each other. Thus, the probabilities of both experiencing good periods (*p*_*GG*_), both experiencing bad periods (*p*_*BB*_), type 1 experiencing good and type 2 experiencing bad (*p*_*GB*_), and vice versa (*p*_*BG*_) are given by:

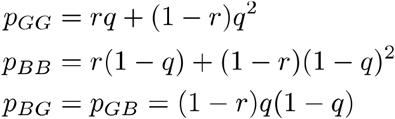

As they must *p*_*GG*_, *p*_*BB*_, *p*_*BG*_ and *p*_*GB*_ sum to 1.

Let *x*_*k*_(*t*) be the number of cells of type *k* in an arrested state following time period *t*, where *k*=1 or *2*. The two species growth equations take on the following form:

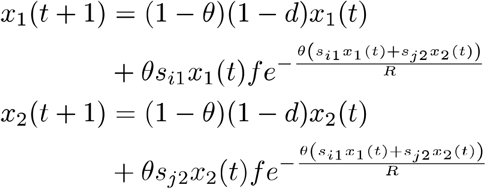

where *i*, *j* = *g* or *b*, and *d* is the mortality rate of cells in an arrested state per period, and *R* scales the limits to growth by representing some measure of resources available to cells that are in a proliferative state.

The above model easily generates a coexistence via a storage effect. This can be seen by showing how either species 1 or 2 can invade a resident population of the other, and that when together they settle on a dynamic equilibrium but with more or less temporal variation in the frequency of the two cell types within a micro-environment (**Fig. 1**). When averaged over many micro-environments the cell type frequencies might appear relatively stable across time. If a storage effect is operating within a tumor, then there may be surprisingly large fluctuations in cell composition at the level of small neighborhoods of cancer cells.

**Figure 1:**
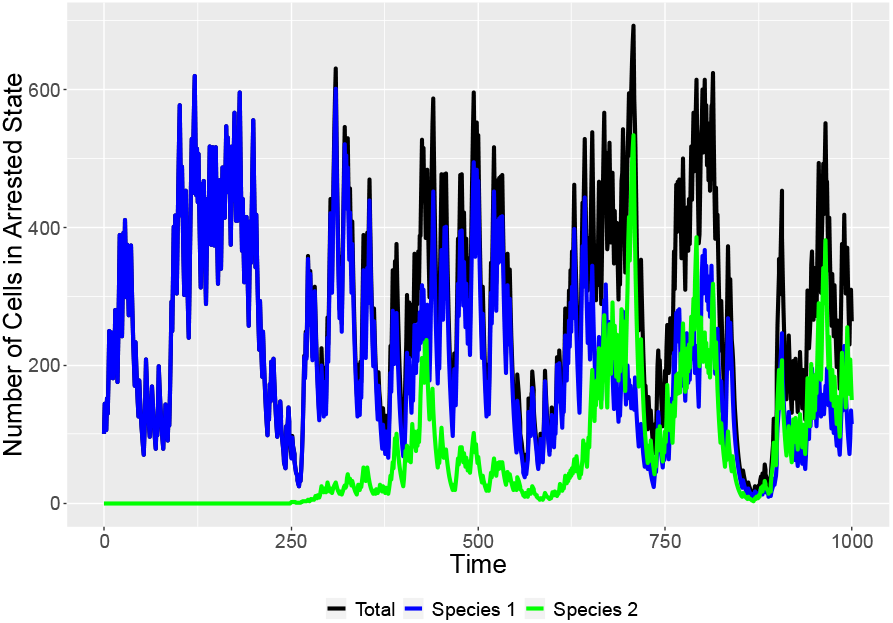
Invasion of resident population. Parameters: *s*_*g*_=0.8, *s*_*b*_=0.1, *R*=100, *d*=0.05, *θ*=0.2, *f*=5, *q*=0.4, r=0.8, *x*_*1*_(0)=100, *x*_*2*_(0)=0, *x*_*2*_(250)=1.

The storage effect is not possible in the absence of an arrested state. For θ = 1, the two cancer cell types exist as a stochastic random walk with neither one having an advantage. An arrested state buffers a cell type when it experiences a bad period while the other experiences a good period. Similarly, if *r* = 1, then a storage effect is not possible, as both cell types perceive good and bad years identically. The two will simply experience a stochastic random walk of population sizes with neither experiencing an advantage when rare. Once *r* < 1 and *θ* < 1, coexistence of cell types by the storage effect will happen, but the strength of this will increase as *r* approaches 0. The advantage that a cell type gains when it is rare (thus driving it to increase in time at the expense of the other cell type) derives from the occurrence of periods where the rare species experiences a good and the resident cell type experiences a bad period, giving the rare species opportunity to realize the high growth potential of a good period without strong depression by competition. This is maximized when *r* = 0 and when q = 0.5 (good and bad periods are equally likely). Furthermore, as a long-term dynamic equilibrium, the average tumor burden increases as *r* goes from 1 to 0; and as *q* goes from 0 to 1 (**Fig. 2**).

**Figure 2:**
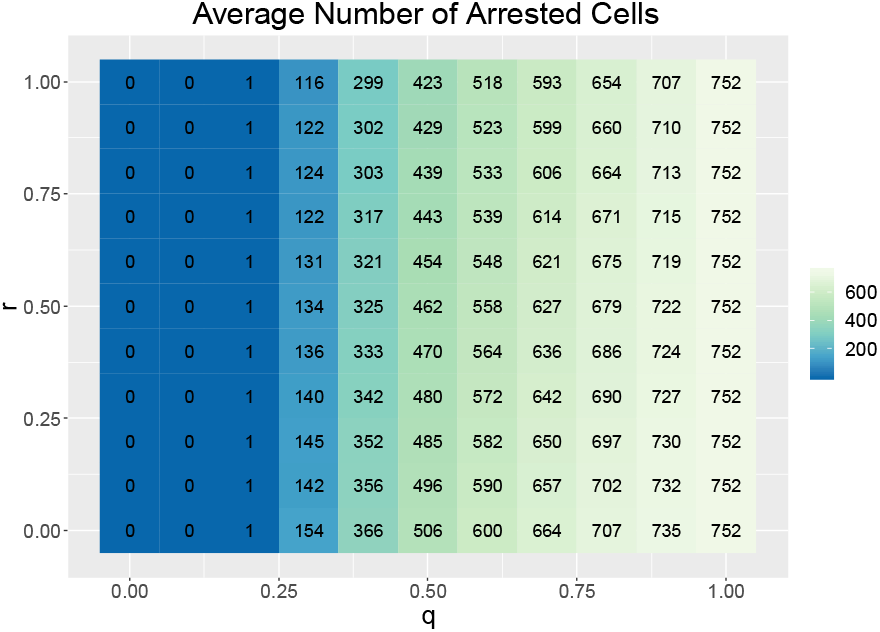
Average number of arrested cells as a function of q and r. Each combination represents an average of 10 simulations. Parameters: *s*_*g*_=0.8, *s*_*b*_=0.1, *R*=100, *d*=0.05, θ=0.2, *f*=5, *x*_*1*_(0)=100, *x*_*2*_(0)=100, number of timesteps=10^4^.

## How Might Consideration of the Storage Effect Inform Cancer Therapy?

The existence of the storage effect contributing to diversity in cancer would have consequences for clinical treatments. While intratumor heterogeneity is established as a marker for poor prognosis, an underlying storage effect dynamic would present new challenges. The presence of a storage effect presents a double challenge for therapy. First is the task of targeting quiescent cells, and second is the challenge of finding drugs effective on each or all of the cancer cell types.

Quiescence or dormancy now appears to be a major mechanism by which cancers evade drug therapies and initiate metastases^23–25^, so there is great interest in whether quiescent or dormant cells can be targeted by therapies^61^. This paper argues that we must also consider that there may be many coexisting clonal lineages of cancer cells that differ in physiologies of their non-proliferative and proliferative life history states. If cancer cell types coexist via the storage effect, then one is not targeting just a single cell type, but a diversity of cell types with potentially differing susceptibilities to therapy.

### Pseudo-resistance

Heterogeneity in quiescence could obscure detection of resistance to treatment through the phenomenon of p*seudo-resistance*, a situation where an otherwise treatment-sensitive population of cancer cells appears to be resistant. Pseudo-resistance could result from temporal variation that selects for cell lineages that maintain a high fraction of cells within a quiescent state. Many cytotoxic chemotherapies target proliferating cells that are actively dividing, and non-proliferative cells may be insensitive to these therapies. Those parts of a tumor that experience the greatest temporal variability may have the greatest fraction of non-proliferative cells. In this case, therapy may appear effective in eliminating tumor cells, but continued efficacy may slow, as the remaining cancer clonal populations have lower fractions of cells in a proliferative state. The storage effect, by maintaining the coexistence of cancer clones that vary in their propensity for remaining quiescent or arrested, could exacerbate pseudo-resistance.

With pseudo-resistance, continuing the same drug regimen would continue to cull cancer cells as they emerge into a proliferative state. However, the fraction of cancer cells killed with each dosing would decline, generating the appearance of declining tumor burden, but without entirely eradicating the tumor. Prolonging the therapy may increase the likelihood of true resistance emerging from the surviving cancer cells and also may induce undue toxicities to the patient. The possibility that pseudo-resistance might reflect a storage effect would suggest novel therapeutic strategies. One approach might be to maintain the ongoing therapy, but reduce dosage or make it more intermittent (akin to maintenance therapy for patients in remission). Frequent and accurate measures of tumor burden might identify breakpoints in the rate of tumor burden decline, which would indicate the presence of different clones with different propensities to remain quiescent. A second therapeutic approach could be to include in tandem an intervention designed to amplify the rate at which cancer cells return from quiescence into a proliferative state. It also might be useful to begin with the first therapy and then add to it, rather than replace it with, additional therapies; if possible, the second therapy would be chosen for toxicity to quiescent cells.

### The Storage Effect and Multi-drug Therapies

Existence of a storage effect implies that additional data is needed to predict tumor progression. Knowing to what extent the variability that one sees in space also occurs in time at each spot becomes a valuable and critical piece of information. There is recognition of the need for serial histologies, but the destructive nature of histological sampling means exact resampling is impossible. Variable temporal dynamics means that local fluctuations could work against traditional therapies. An underlying storage effect would indicate that cell types differ in their dependence on or sensitivity to specific micro-environmental conditions that are fluctuating. Each of them could have an Achilles’ heel. If one can identify or suggest the tradeoff promoting a storage effect, then a background therapy targeting quiescent cells could be combined with a standard chemotherapy and a therapy known to target the specific attributes of the cancer cell types.

## Summary

Here, we have described the diversity-promoting mechanism known as the storage effect and suggested that the conditions that make the storage effect relevant to ecosystems in nature also are found in cancer. Research is needed to better identify and quantify the features of cancer that characterize the storage effect. These features include population-specific responses to the environment, which could give differential use of conditions in time by different cancer cell populations; and behaviors associated with entry into, exit from, and duration of quiescence in cancer cells, which could enable buffering of those populations to environmental fluctuations, allowing them to evade losses in bad times and realize strong growth in good times. The possibility that the storage effect is involved in the clonal ecological heterogeneity and quiescence/dormancy of cancer cells, both of which pose serious conundrums for cancer therapies, dictates that more attention should be paid to understanding the temporal dimensions of cancer. To do this, both the data and the models that inform our understanding of cancer must be expanded to better reveal and account for temporal variation. If temporal variation in environmental suitability for growth contributes significantly to population interactions within the cancer ecosystem, the implications to how we understand and treat cancer would be substantial.

## Acknowledgements

The authors would like to thank Drs. Sarah Amend, Ryan Bishop, and Chris Whelan for helpful discussions on the content of this paper.

